# Neural Encoding of Direction and Distance across Reference Frames in Visually Guided Reaching

**DOI:** 10.1101/2024.09.19.613668

**Authors:** Alejandra Harris Caceres, Deborah A. Barany, Neil M. Dundon, Jolinda Smith, Michelle Marneweck

## Abstract

Goal-directed actions require transforming sensory information into motor plans defined across multiple parameters and reference frames. Substantial evidence supports the encoding of target direction in gaze– and body-centered coordinates within parietal and premotor regions. However, how the brain encodes the equally critical parameter of target distance remains less understood. Here, using Bayesian pattern component modeling of fMRI data during a delayed reach-to-target task, we dissociated the neural encoding of both target direction and the relative distances between target, gaze, and hand at early and late stages of motor planning. This approach revealed independent representations of direction and distance along the human dorsomedial reach pathway. During early planning, most premotor and superior parietal areas encoded a target’s distance in single or multiple reference frames and encoded its direction. In contrast, distance encoding was magnified in gaze– and body-centric reference frames during late planning. These results emphasize a flexible and efficient human central nervous system that achieves goals by remapping sensory information related to multiple parameters, such as distance and direction, in the same brain areas.

**Significance statement:** Motor plans specify various parameters, e.g., target direction and distance, each of which can be defined in multiple reference frames relative to gaze, limb, or head. Combining fMRI, a delayed reach-to-target task, and Bayesian pattern component modeling, we present evidence for independent goal-relevant representations of direction and distance in multiple reference frames across early and late planning along the dorsomedial reach pathway. Initially, areas encoding distance also encode direction, but later in planning, distance encoding in multiple reference frames was magnified. These results emphasize central nervous system flexibility in transforming movement parameters in multiple reference frames crucial for successful goal-directed actions and have important implications for brain-computer interface technology advances with sensory integration.

## Introduction

Visual goal-directed reaching relies on neuronal events that transform visual information about a target location into an actionable plan to move the chosen effector (Crawford et al., 2011; Fooken et al., 2023). Achieving a motor goal necessitates specifying not just the direction, but critically also the distance to the target. Complexity arises because parameters can be specified in multiple reference frames relative to the gaze, head, or limb (Pouget et al., 2002; Battaglia-Mayer et al., 2003; Fiehler and Karimpur, 2023). While direction has been extensively studied, how the brain encodes distance remains less understood. Determining how the brain supports all manner of sensorimotor transformations is an urgent challenge considering the gains in robotic control with brain-computer interface technology integrating sensory information (Flesher et al., 2021).

A substantial body of research has demonstrated that encoding movement parameters in multiple coordinate frames provides the flexibility and efficiency needed for optimal sensorimotor transformations, however, this has been primarily studied in relation to target direction. Monkey electrophysiology studies and human functional neuroimaging, neurostimulation, and lesion studies have identified the superior parietal cortex (SPL) and dorsal premotor cortex (PMd) as critical nodes for computing reaching actions (Wise and Mauritz, 1985; Kalaska and Crammond, 1995; Galletti et al., 1997; Graziano and Gross, 1998; Crammond and Kalaska, 2000; Andersen and Buneo, 2002; Bastian et al., 2003; Johnson and Grafton, 2003; Rizzolatti and Matelli, 2003; Gallivan et al., 2009; Pisella et al., 2009; Fabbri et al., 2010; Kaufman et al., 2010; Gallivan et al., 2011a; Gallivan et al., 2011b; Davare et al., 2012; Barany et al., 2014; Fabbri et al., 2014; Kaufman et al., 2014; Coallier et al., 2015). SPL and PMd encode a target’s direction in body and gaze-centric reference frames often via overlapping neuronal populations within this dorsomedial reach stream (Boussaoud et al., 1998; Buneo et al., 2002; Batista et al., 2007; Marzocchi et al., 2008; Bernier and Grafton, 2010; Beurze et al., 2010; Chang and Snyder, 2010; Bremner and Andersen, 2012; Hadjidimitrakis et al., 2014a; Leoné et al., 2015; Bosco et al., 2016; Piserchia et al., 2016; Cappadocia et al., 2017; De Vitis et al., 2019; Magri et al., 2019). Encoding target direction in multiple frames supports computational models that emphasize the flexibility, efficiency, and power of individual nodes to optimally transform between extrinsic or intrinsic inputs (Körding and Wolpert, 2004; Sober and Sabes, 2005; McGuire and Sabes, 2009).

Despite its equally critical role in successful reaching, much less is known about how the brain encodes target distance. Behavioral studies suggest that direction and distance are specified independently (Gordon et al., 1994; Messier and Kalaska, 1997; Vindras et al., 2005). There is a growing number of datasets on non-human primates showing distance, like direction, is encoded in multiple reference frames (Hadjidimitrakis et al., 2014a; Bosco et al., 2016; Piserchia et al., 2016; Hadjidimitrakis et al., 2017; De Vitis et al., 2019; Hadjidimitrakis et al., 2020) and processed after direction in PMd (Messier and Kalaska, 2000; Churchland et al., 2006; Davare et al., 2015) and SPL (Hadjidimitrakis et al., 2014b; Hadjidimitrakis et al., 2022). In humans, transcranial magnetic stimulation of SPL before movement onset resulted in directional errors, whereas PMd stimulation at a slightly later time point disrupted accurate computations of movement distance, supporting serial processing of direction, then distance (Davare et al., 2015). During movement execution, SPL and PMd represent both distance and direction features (Fabbri et al., 2012). These similarities—including independent specification, serial processing, neural overlap, and shared susceptibility to disruption—suggest that distance, like direction, may also have a flexible representation across multiple reference frames. However, the coordinate frame(s) in which distance is planned is unknown, especially in humans.

The primary aim of this study was to determine the reference frame in which a target’s distance is represented in the human dorsomedial reach pathway during early and late planning of an upcoming motor action. One hypothesis is that, unlike direction, a target’s distance is encoded strictly within a single reference frame, reflecting a hierarchical cost-efficient strategy where only some movement parameters (i.e., direction) but not all (i.e., distance) undergo sensorimotor transformations (Turella et al., 2020). Alternatively, distance might be encoded in multiple reference frames, similar to direction (Bosco et al., 2016), which suggests a similar circuitry that flexibly transforms between gaze and body-centered coordinate spaces for distance as it does for direction. We also aimed to investigate whether distance representations become more pronounced during later stages of planning, in line with serial processing accounts.

To address these questions, we employed fMRI with a delayed reach-to-target task, varying the distance (near vs. far) between target and gaze, target and hand, and gaze and hand, as well as the direction (left vs. right) of targets, gaze, and hand position (Fig. 1). Using variational representational analysis (vRSA) implementing Bayesian pattern component modeling of fMRI data (Friston et al., 2019; Marneweck et al., 2023; Kreter et al., 2024), we determined how distance and direction parameters are represented in SPL, PMd, and primary motor cortex (M1) during early and late planning. Our results show that during early planning, some regions encoded target distance in a single reference frame, mostly gaze-centric, while other areas represented this information in multiple reference frames. Notably, all regions encoding distance information also encoded direction information independently. During late planning, consistent with serial processing accounts, distance representational specificity in multiple reference frames magnified across the dorsomedial reach pathway. These findings demonstrate how and when multiple goal-relevant representations are specified in humans, supporting the hypothesis that similar circuitry flexibly transforms between gaze and body-centered coordinate spaces for distance as it does for direction.

**Figure 1.**
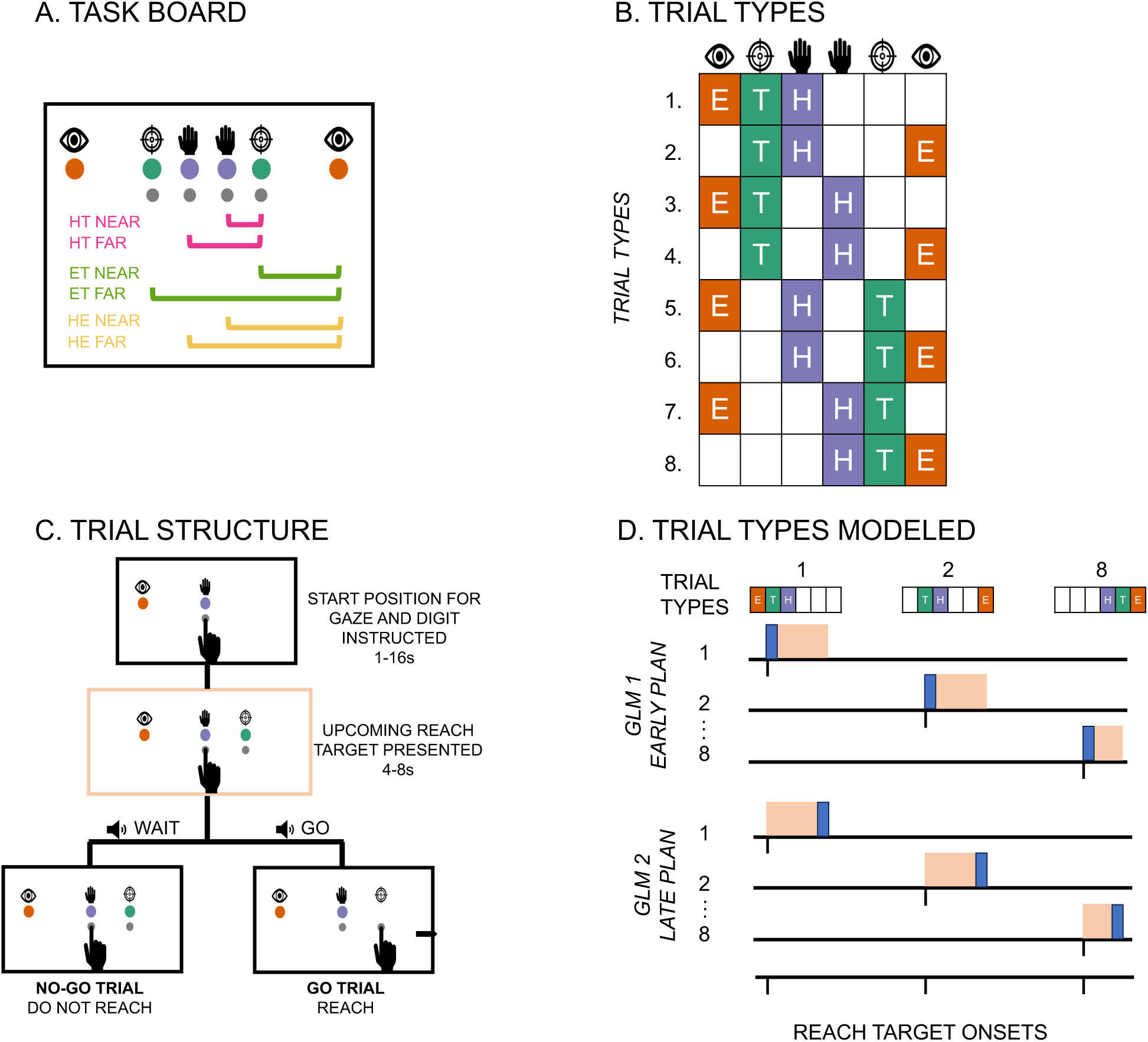
**A**. Schematic of the reach-to-target task board. Colored LEDs and icons showed hand, target, and gaze positions, and two buttons for hand and target positions, respectively. Near and far vector distances between target and gaze (eye-target), target and hand (hand-target), and between hand and gaze (hand-eye) are visualized but not present on the actual board. **B**. Eight task trial types with gaze (E), initial hand (H), and target (T) position on the left or right side of the board gave two possible target, gaze and hand position directions (left vs. right) and two possible distances (near vs. far) between target and gaze, target and hand, and gaze and hand. **C.** Each trial started with a hand and gaze position instructed, followed by the appearance of a goal target, after which an audio cue instructed a go or a no-go trial. No-go trials were used for analyses to prevent movement execution contamination in early and late plan-related BOLD activity. **D.** General linear model (GLM) structures where each GLM models the extent to which a trial type predicts BOLD activity during either the early (first 2-sec) or late planning (last 2-sec) period during which an upcoming target is presented. The resultant beta-estimates then serve as inputs to variational representational analysis (vRSA) of fMRI data to test the extent to which a region of interest represents features or components such as a target, gaze, or hand direction, or the distance between each of these during early and late planning.

## Materials and Methods

### Participants

Twenty-seven right-handed young adults with normal or corrected-to-normal vision participated in this study (14 female, *M* = 21.1 ± 3.3 years). All participants reported having no history of neurological or neuromuscular diagnoses, and no motor or cognitive impairments that would adversely affect their ability to perform motor tasks. All participants gave written informed consent, and the experimental procedures were approved by the University of Oregon Institutional Review Board.

### Materials, design, and procedure

#### Materials

An interactive task board was held by a stand 4 cm above the table and angled 40.6° relative to the table, which piloting showed as an optimal orientation for full view. As shown in Fig. 1A, the task board base (width: 40.0 cm; length: 10.0 cm; depth: 6.5 cm) consisted of six colored LEDs that, when illuminated, would instruct a left-or right-sided gaze position (red), initial hand position (green), and target position (blue), respectively (we ensured all participants could discriminate between these colors before scanning). Four buttons to be pressed when instructed were positioned below each of the hand and target position LEDs. Icons illustrating the eye, hand, and target position were placed above each of the LEDs. The two hand LEDs were 2.3 cm apart and positioned in the middle of the board. On each of the outer sides of these hand LEDs 2.3 cm away were two target LEDs positioned 6.9 cm apart from each other. The gaze LEDs were 32.9 cm apart and positioned on the outer sides of the target LEDs.

The configuration of gaze, hand, and target position on the task board gave two possible distances between gaze and target (ET near: 13.0 cm; ET far: 19.9 cm), and hand and target (HT near: 2.3 cm; HT far: 4.6 cm). Visual angles between gaze position and targets for ET-near and ET-far trials were 11.3° and 16.8°, respectively (from the distance of the bridge of the nose to the center of task board lights based on data from CDC-defined average height of females and males). Distances were selected to minimize visual acuity differences between ET near and far such that targets in both cases fell within an observer’s 5-20° near-peripheral zone within which there are little changes in the parameters of acuity (Millodot et al., 1975; Larson and Loschky, 2009).

Eye tracking was performed outside the scanner with Pupil Core glasses and the head secured in a headrest, with an eye camera resolution and frequency of 192 x 192 pixels and 200 Hz, and a scene camera resolution and frame rate of 1280 x 720 and 30 Hz (Kassner et al., 2014).

Anatomical and fMRI data were collected using a Siemens 3T Skyra (32-channel phased-array head coil). High-resolution 1 mm isotropic T1-weighted (TR: 2500 ms, TE: 2.98 ms, FA: 7°, FOV: 256 mm) and T2*-weighted (TR: 3200 ms, TE: 565 ms, FOV: 256 mm) images were acquired of the whole brain. Both sequences used volumetric navigators for prospective motion correction and selective reacquisition (Casey et al., 2018). Next, as subjects performed the reach-to-target task, BOLD contrast was measured with a multiband T2*-weighted echoplanar gradient echo imaging sequence (TR: 450 ms, TE: 30 ms, FA: 45°, FOV: 192 mm, multiband factor: 6). Each functional image consisted of 48 slices acquired parallel to the AC–PC plane (3 mm thick; 3 x 3 mm in-plane resolution) (Moeller et al., 2010; Setsompop et al., 2012).

#### Experimental design and procedure

During fMRI, participants performed the reach-to-target task with their right hand on a task board at arm’s length sitting on a marked spot on a table positioned over the hips of the subject. With a mirror attached to the head coil, subjects saw the task board and their hand (statically and during movement) as if they looked directly at it when sitting upright.

Participants reached to left-or right-sided targets, with initial hand position and gaze fixated in one of two locations, respectively (Fig. 1A, B). As seen in Fig. 1C, each trial began with illuminated LEDs that instructed an initial hand position and a gaze position to focus on for the entire trial, which piloting showed were successfully adhered to (see below, Table 1). After a variable delay period (1, 2, 4, 8, 16-s with the respective proportions of trials, [0.52, 0.26, 0.13, 0.06, 0.03]), a target LED illuminated for 4, 6, or 8-s (with the respective proportions of trials [0.56, 0.30, 0.14]). Next, an audio cue played through headphones indicating whether to perform the planned reach to target (*i.e.*, “go”, 60% of the trials) or not (*i.e.*, “wait”, 40% of the trials). The initial hand position and target position LEDs were turned off concurrently with the go/no-go cue to encourage planning the reach during this variable delay period during which the target was illuminated (Ariani et al., 2022). Feedback audio cues indicating success or error played after a correct or incorrect reach to the target, or 2.5-s after the go/no-go cue if no movement was made with an additional 1-s interval between the feedback cue and the start of the next trial. There were eight trial types with two possible initial hand, target, and gaze positions, with the vector distance between the gaze and target, hand and target, and hand and eye varied in one of two ways (near vs. far) (Fig. 1B). After standardized instructions and 20 practice trials, participants completed six functional runs of 40 trials (five trials for each of the eight trial types). Stimulus and button press timings for each trial were controlled by a custom Python script.

**Table 1.** Percent time, in a given trial in maintaining gaze within a predefined surrounding boundary organized by each combination of left-(L) and right-sided (R) gaze (E), initial hand (H), and target (T) position in two participants.

| *H-T-E* | *L-R-L* | *L-R-R* | *L-L-R* | *R-R-R* | *R-L-R* | *R-L-L* | *L-L-L* | *R-R-L* |
| --- | --- | --- | --- | --- | --- | --- | --- | --- |
| *Subj. 1* | 97.4 | 100.0 | 100.0 | 99.5 | 94.5 | 100.0 | 99.2 | 100 |
| *Subj. 2* | 99.6 | 99.4 | 99.8 | 98.8 | 100.0 | 99.6 | 100.0 | 100.0 |

The order and distribution of each of the eight trial types, go and no-go trials, and within-trial variable durations (delay and plan phases) was determined with the goal of minimizing the variance inflation factor (VIF) for each functional run (Ariani et al., 2022)); VIF= var(E)/ var(X), where var(E) is the mean estimation variance of all the regressor weights and var(X) the mean estimation variance had these regressors been estimated in isolation). VIF estimates the severity of multicollinearity between model regressors by providing an index of how much the variance of an estimated regression coefficient is increased because of collinearity. Large VIF values indicate regressors are not independent of each other, whereas a VIF of 1 means no inflation of variance between regressors. We optimized the design such that the VIF was below 1.15, indicating the independence of regressors. In addition, no evidence of multicollinearity was confirmed post-data collection.

#### Hand and Eye Movement Tracking

Reaction time and reach velocity were calculated for every go trial during scanning. Reaction time was calculated as the difference between the time from the go cue to when the hand released the initial hand position button. Reach velocity was calculated as the time from the initial hand position button release to a target button press given the distance traveled between the initial hand position and the target position. Pilot eye-tracking data (*n* = 2) outside the MR with the head secured in a headrest during a 40-trial run showed high task adherence to gaze instruction (Table 1). Pupil Player software (v3.5.1) calculated gaze accuracy, defined as the mean distance difference in the visual angle of the fixation location and instructed gaze location. Mean gaze estimation is reported with an accuracy of 0.6 degrees with 0.08 degrees of precision (Kassner et al., 2014). Raw data was filtered to exclude blinks and recorded values with a confidence level below 0.6. Both subjects maintained gaze within a 3.5 cm x 3.5 cm boundary surrounding each illuminated gaze LED for each trial, in all combinations of gaze, target, and hand positions (*p* > 0.05, Table 1).

#### MRI preprocessing and statistical analyses

Functional images across runs were spatially realigned to a mean EPI image using a second-degree B-spline interpolation, coregistered to each individual subject’s T1, and normalized between-subjects using SPM (<u>fil.ion.ucl.ac.uk/spm</u>). Head motion mean rotations and translations (with minimum and maximum values in parentheses) were minimal: x: 0.02 mm (–2.2, 2.02); y: 0.1 mm (–1.6, 3.3); z: 0.2 mm (–3.9, 4.2); pitch: –0.001 (–0.2,; roll: 0.0008 (–0.04, 0.03); yaw: 0.0008 (–0.04, 0.04).

Following preprocessing of fMRI data, we used vRSA (Friston et al., 2019; Marneweck and Grafton, 2020c, b, a; Marneweck et al., 2023; Kreter et al., 2024) with an adaptation to the Matlab script DEMO_CVA_RSA.m available in SPM12. First, we estimated convolution-based GLMs in SPM12 for each subject and functional run separately with the Robust WLS Toolbox selected to down weight volumes with high noise variance to account for movement artifact (Diedrichsen & Shadmehr, 2005). We generated GLMs that estimated the extent to which planning a reach to target predicted BOLD activity for each of the eight trial types on no-go trials and on go-trials (Fig. 1D). Planning was modeled during 2-s epochs with onsets marked at target presentation (i.e., early plan epoch) and during the last 2-s before the go/no-go cue (i.e., late plane epoch). Similar to several recent fMRI studies on motor planning, our analyses focused on no-go trials to prevent contamination of movement execution in GLM-derived estimates of activity during the planning phase (Ariani et al., 2018; Ariani et al., 2022; Yewbrey et al., 2023). Planning of go trials, movement on each trial, and error trials (if any were made) were modeled as regressors of no interest and not used in further analyses.

Following the GLM step, we extracted GLM-derived beta values in primary motor, dorsal premotor, and superior parietal regions (Fig. 2) from the Julich Brain Atlas (Amunts et al., 2020) and from the Human Brainnetome Atlas (Fan et al., 2016) using FSL’s fslmeants for each participant and functional run. The dorsal premotor area in the Julich atlas is subdivided cytoarchitectonically into three regions (6d1, 6d2, 6d3) in a rostro-caudal arrangement: 6d1 is the most caudal area, located on the precentral gyrus and a caudal part of the superior frontal gyrus (analogous to Glasser atlas area 6d) (Glasser et al., 2016). 6d2 is located rostrally to 6d1 on the superior frontal gyrus (analogous to Glasser atlas area 6a), and 6d3 is located exclusively in the superior central sulcus, mainly ventrally to 6d2 (analogous to Glasser atlas area 6a). Hereafter we refer to 6d1 as PMd caudal, 6d2 as PMd rostral, and 6d3 as PMd rostro-sulcal. The location and extent of key superior parietal areas correspond with the same areas as that in the Glasser atlas (Glasser et al., 2016), including SPL 7a and SPL 5m. Additionally, Julich area SPL 7a comprises mIPS per the Glasser atlas and previous human neuroimaging work (Bernier and Grafton, 2010) and pIPS per Fabbri et al. (2012).

**Figure 2.**
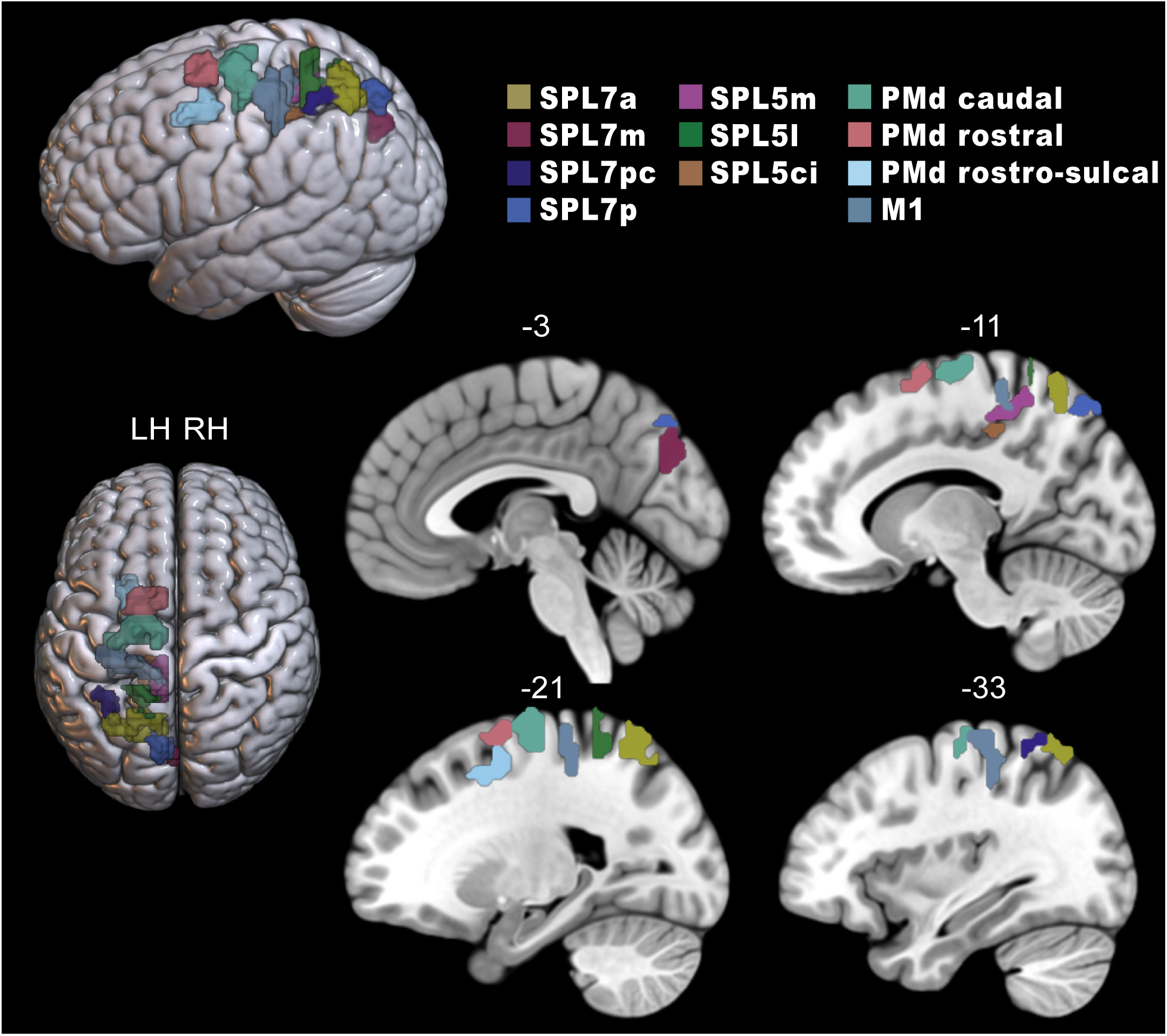
Predefined regions of interest extracted from the Julich Brain Atlas and Human Brainnetome atlas (primary motor cortex, M1) are displayed on the MNI-152 atlas using the visualization software, MRIcroGL. SPL: superior parietal; PMd: dorsal premotor.

We populated a condition (*i.e.,* eight)-by-voxel U matrix describing each condition’s voxel activity pattern (calculated with the above GLM) from which we computed a condition-by-condition second-order similarity matrix (G) that describes the relationship between these activity patterns (*i.e.*, 8 x 8 covariance matrix), where G=UU^T^. Higher values in off-diagonal cells reflect higher pattern similarity between conditions.

Next, we tested hypotheses regarding the composition of these second-order statistics by inferring the contribution of ‘components’ to the G covariance matrix. Fig. 3 shows the components we included. Our main components of interest test the extent to which a region represents a target’s distance to the hand and gaze, respectively, during early and late planning (Fig. 3A). We also include direction-specific components that varied in the task design, including target position (absolute or relative to hand/body), initial hand position (absolute or relative to body), gaze positions (absolute or relative to hand/body/target), and the hand-gaze distance, in each model (Fig. 3B). A strength of the vRSA approach is to test for the independent contribution of each component to the covariance G matrix (similar to multiple regression). The method quantifies the contribution of multiple components while taking into account all specific contrasts. In this way, an ROI can be sensitive to more than one component (*i.e.*, such that a region’s spatial activity patterns can represent multiple conditions of interest). Moreover, the covariance G matrix does not depend on the way columns of U are ordered (similar to a regression coefficient that will not change if shuffling the order of contributing pairwise x-y data). Pattern similarity can be identified even if not every column contributes to the covariance matrix.

**Figure 3.**
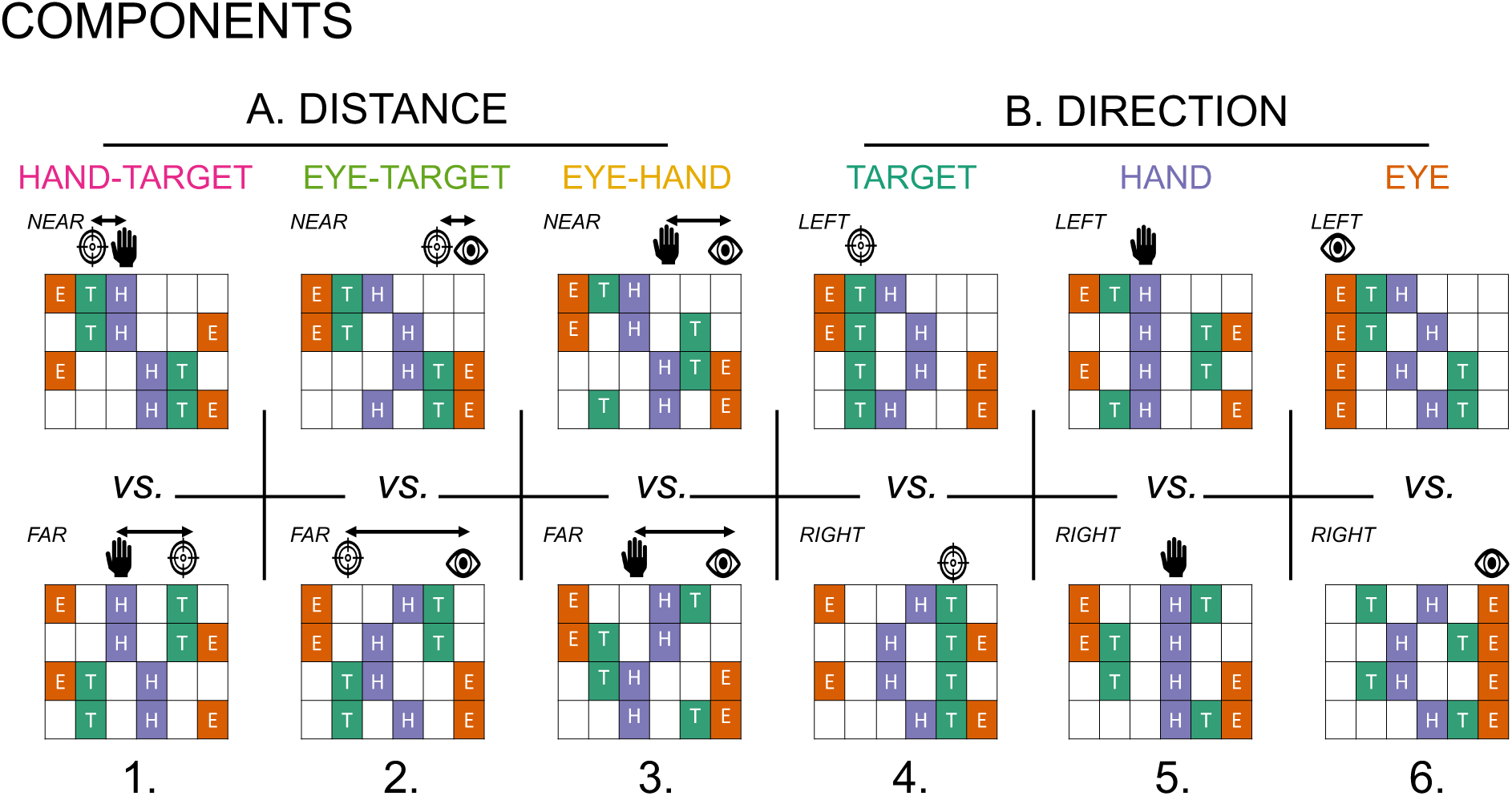
Variational representational similarity analysis (vRSA) implements Bayesian pattern component modeling of fMRI data tested for component effects of (A) the distance between 1) target and hand (HT), 2) target and eye (ET), and 3) hand and eye (HE), and (B) the direction of a 4) target position (T), 5) initial hand position (H), and 6) eye position (E).

Within this pattern component modeling framework, we could simultaneously test if and how distance between target, gaze, and hand position are represented, after accounting for direction components known to be encoded throughout this network of regions (Wise and Mauritz, 1985; Kalaska and Crammond, 1995; Galletti et al., 1997; Graziano and Gross, 1998; Crammond and Kalaska, 2000; Andersen and Buneo, 2002; Bastian et al., 2003; Johnson and Grafton, 2003; Rizzolatti and Matelli, 2003; Gallivan et al., 2009; Pisella et al., 2009; Fabbri et al., 2010; Kaufman et al., 2010; Gallivan et al., 2011a; Gallivan et al., 2011b; Davare et al., 2012; Barany et al., 2014; Fabbri et al., 2014; Kaufman et al., 2014; Coallier et al., 2015). Evidence of distance-based representations would be indicated by distinct activity between near vs. far targets relative to 1) the initial hand position (Hand-Target, HT), 2) the instructed gaze position (Eye-Target, ET), and between near vs. far distances between the initial hand position and the instructed gaze position (Hand-Eye, HE). Evidence of direction-based representations would be indicated by distinct activity patterns of left-vs. right-sided targets, initial hand positions, and gaze positions. We report results from vRSA’s run with GLM data modeling the 2-s at the start and the end of the planning epoch, respectively.

vRSA performed with SPM functions returns log evidence (marginal evidence) enumerating each component’s contribution to the second-order matrix (G) for a given ROI at the group level, where more negative values are greater evidence of a component’s contribution. To establish a criterion that sufficient evidence is observed, we take an ultra-conservative step, detailed in Marneweck et al., 2023, by comparing actual log evidence with a null distribution of log evidence values for each component in each ROI. We derive each null distribution of log evidence by shuffling condition labels (note – not voxel order) in our U matrix 1000 times, each time estimating G from the shuffled data and enumerating the component contributions. We can then illustrate the log evidence values of our real data (*i.e.*, unshuffled) relative to these null distributions. We subtract the log evidence of the correct condition label from each of the shuffled log evidence values, creating a distribution of log Bayes factors, where higher values now communicate a strong effect. We conclude strongly credible effects only if there is little to no overlap between the null distribution and model evidence. We then formalize strong evidence for a component to contribute to a region’s activity pattern if the real Bayes factor is three times more credible than the 80% or 95% strongest effect from the null distribution. See Marneweck et al. 2023 for a comprehensive discussion on how this conservative approach mitigates the need for multiple comparison corrections within and across ROIs (also see (Gelman et al., 2012; Gelman & Loken, 2016; Krushke, 2015; Marneweck & Grafton, 2020)).

## Results

During fMRI, participants performed a delayed reach-to-target task, adapted from similar tasks used in nonhuman primates (Batista et al., 1999; Buneo et al., 2002; Pesaran et al., 2006; Batista et al., 2007), with two possible distances (near vs. far) between hand and target, gaze and target, and gaze and hand, and two possible directions (left vs. right) for target, gaze and hand positions, respectively. The primary aim of this study was to determine if and how the distance of an upcoming reach target is represented in SPL, PMd, and M1 regions (Fig. 2) during early and late planning epochs. Secondarily, we determined if distance representations are magnified during later planning. Our analyses focused on plan periods of no-go trials to avoid movement execution contaminating plan-related BOLD activity (Ariani et al., 2022; Yewbrey et al., 2023).

Using vRSA of fMRI data (Friston et al., 2019; Marneweck and Grafton, 2020a, b, c; Marneweck et al., 2023) in predefined M1, PMd, and SPL ROIs, we determined the extent to which components of interest modulate overall condition-by-condition differences in neural activity patterns. We focused on six distinct components: distance between 1) hand and target, 2) gaze and target, and 3) hand and eye, and the direction of a 4) target, 5) initial hand position, and 6) gaze position (Fig. 3). The strength of the vRSA approach is that it tests for the independent contribution of each component to the condition-by-condition covariance matrix (similar to multiple regression). In this way, an ROI can be sensitive to more than one component (*i.e.*, such that a region’s spatial activity patterns can represent multiple conditions of interest).

### Distance representations in single or multiple reference frames overlap with direction representations during early planning

We used vRSA to test for each hypothesized component’s contribution to explaining the estimated spatial activity patterns for each condition and ROI during the first 2 s of planning. The vRSA returns log evidence enumerating the strength of each component’s contribution to an ROI’s activation pattern at the group level, with higher values indicating stronger effects (see Methods and Materials for details).

As seen in Fig. 4, strong evidence of representations of a target’s distance were present in multiple parietal (SPL 7a, 7pc, 5m, 5ci), dorsal premotor (PMd caudal, rostral, and rostral-sulcal), and M1 regions. Whereas SPL 7a, PMd rostral, and rostral-sulcal areas encoded target distance in both body and gaze-centric reference frames (HT, ET), other regions encoded target distance in a single gaze-centric (M1, PMd caudal, SPL 5ci, 5m, 7pc) or body-centric reference frame (SPL 5m) during the early stages of planning. Most regions with activity patterns predicted by a target’s distance also contained activity patterns predicted by target, hand, and eye position direction-based components (T, H, and E), and the distance between hand and eye position (HE). This suggests overlap in neuronal populations in SPL and PMd regions that encode direction and distance features critical for an upcoming goal-oriented reach. In addition, some parietal regions represented direction components without distance features (SPL 7m, 7p, 5l), with effects for eye position, indicating widespread direction-based effects. Overall, representational specificity for direction components dominated over that for distance during this early planning epoch, with far more credible effects for direction-based (*n* = 27) than distance-based (*n* = 17) components.

**Figure 4.**
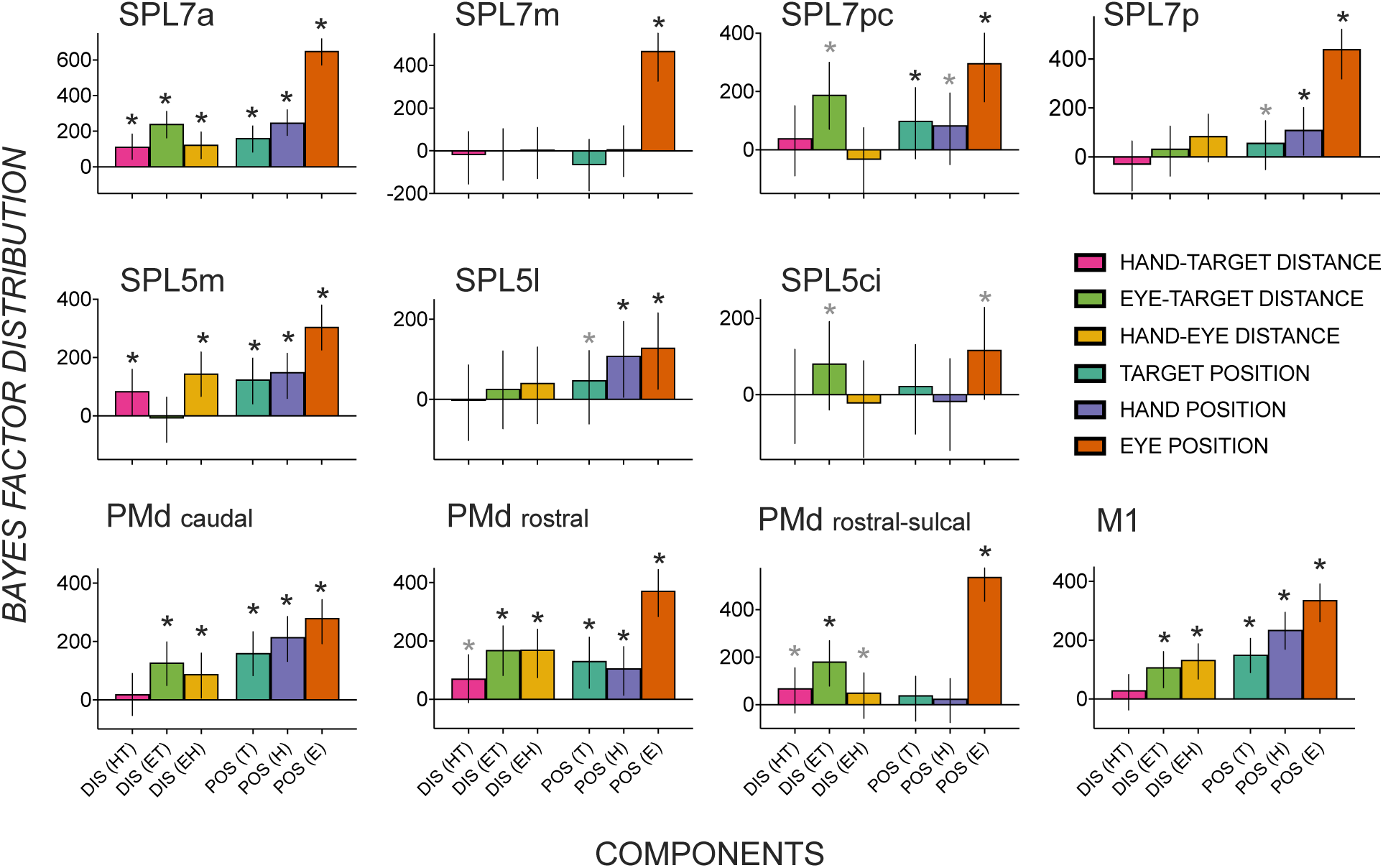
Distribution of Bayes factors (median and highest density interval as error bars) quantifying evidence for spatial activity patterns in dorsomedial reach regions during early planning to be predicted by an upcoming target’s 1) body-centric distance – near vs. far targets relative to the hand (HT); 2) gaze-centric distance – near vs. far targets relative to the gaze (ET); 3) gaze-hand distance (EH) 4) target position – left vs. right (T), 5) initial hand position – left vs. right (H), 6) gaze position – left vs. right (E). Components 1-3 test for distance-related effects, and components 3-6 test for direction-related effects. Asterisk (*) depicts substantial evidence for a component to contribute to a region’s spatial activity pattern that is three times more credible than the 95% (black) or 80% (gray) strongest effect from a null distribution (i.e., shuffled condition labels). SPL: superior parietal; PMd: dorsal premotor; M1: primary motor; DIS: distance; POS: position.

### Heightened representational specificity of a target’s distance during late planning

Fig. 5 shows median Bayes factors quantifying the extent of credible evidence in each ROI for distance and direction-based effects during late planning. A striking observation is a growing number of ROIs encoding the target’s distance in multiple reference frames alongside the relative dissipation of evidence of direction-based activity patterns. Representations of a target’s distance in multiple gaze– and body-centered reference frames in early planning persist in late planning (SPL 7a, PMd rostral, PMd rostral-sulcal). Other regions previously shown to represent a target’s distance in a single reference frame during early planning show evidence of body and gaze-centric encoding during late planning (SPL 5m, PMd caudal, M1).

**Figure 5.**
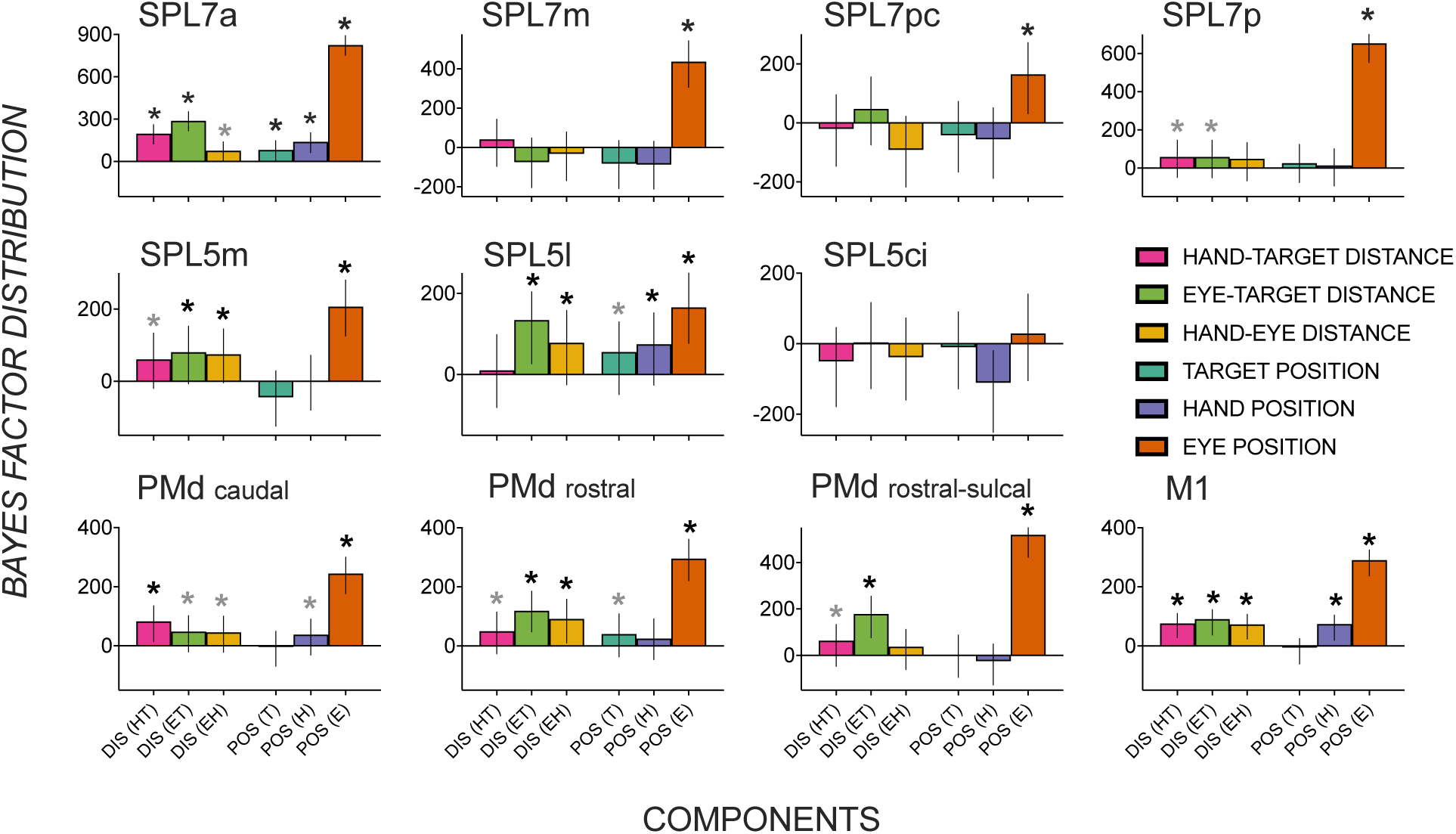
Distribution of Bayes factors (median and highest density interval as error bars) quantifying evidence for spatial activity patterns in dorsomedial reach regions during late planning to be predicted by an upcoming target’s 1) body-centric distance – near vs. far targets relative to the hand (HT); 2) gaze-centric distance – near vs. far targets relative to the eye (ET); 3) eye-hand distance (EH) 4) target position – left vs. right (T), 5) initial hand position – left vs. right (H), 6) eye position – left vs. right – (E). Asterisk (*) depicts substantial evidence for a component to contribute to a region’s spatial activity pattern that is three times more credible than the 95% (black) or 80% (gray) strongest effect from a null distribution (i.e., shuffled condition labels). SPL: superior parietal; PMd: dorsal premotor; M1: primary motor; DIS: distance; POS: position.

Surprisingly, direction-based representations were not as strong as in the early planning period. Multiple regions represented a target position or its direction relative to the hand/body in the early but not in the late phase of planning (SPL 7pc, 7p, 5m, PMd caudal, M1). Similarly, direction-based hand position representations observed during early planning were no longer credibly evident during late planning (SPL 7pc, 7p, 5m, PMd rostral). In contrast to direction-dominant encoding during early planning, credible effects consistent with magnified distance-representational specificity (*n* = 21) marginally exceeded direction-representational specificity (*n* = 17) across dorsomedial reach nodes.

Although direction-based representations relevant to an upcoming reach dissipated in late planning across the frontoparietal network, they were not completely absent. Target position representations persisted in SPL 7a, 5l, PMd rostral. Hand position representations were observed in SPL 7a, 5l, PMd caudal, and M1, and gaze position representations were seen in nearly all tested regions (except SPL 5ci).

### Comparing go and no-go trial activity patterns

The go/no-go approach we adopted here has become an increasingly popular choice for isolating motor planning from execution in fMRI data (Cross et al., 2007; Ariani et al., 2018; Ariani et al., 2022; Yewbrey et al., 2023). We nonetheless briefly report on spatial activity pattern differences between go and no-go trials in key regions revealed by our analyses of the planning period of no-go trials (SPL 7a, 5m, PMd caudal, rostral, rostral-sulcal, and M1). During early planning, we found no differences between go and no-go trials in any of these regions. During late planning, we only found differences in two motor regions (caudal PMd and M1) consistent with previous work (Denyer et al., 2022).

### Activation pattern differences are not explained by behavioral differences

Fig. 6 shows no significant differences in reaction time in initiating a reach and no differences in velocity in reaching for targets, with no significant main effects of components (gaze, hand, target, HT, ET, HE), contrasts (left vs. right, near vs. far), and no interactions (*p*’s > .05). These behavioral measures were taken on go trials. However, they suggest that activity pattern differences reflecting representations of direction and distance during preparatory stages on a no-go trial found in our vRSA results are not predominantly a function of reaction time or velocity differences.

**Figure 6.**
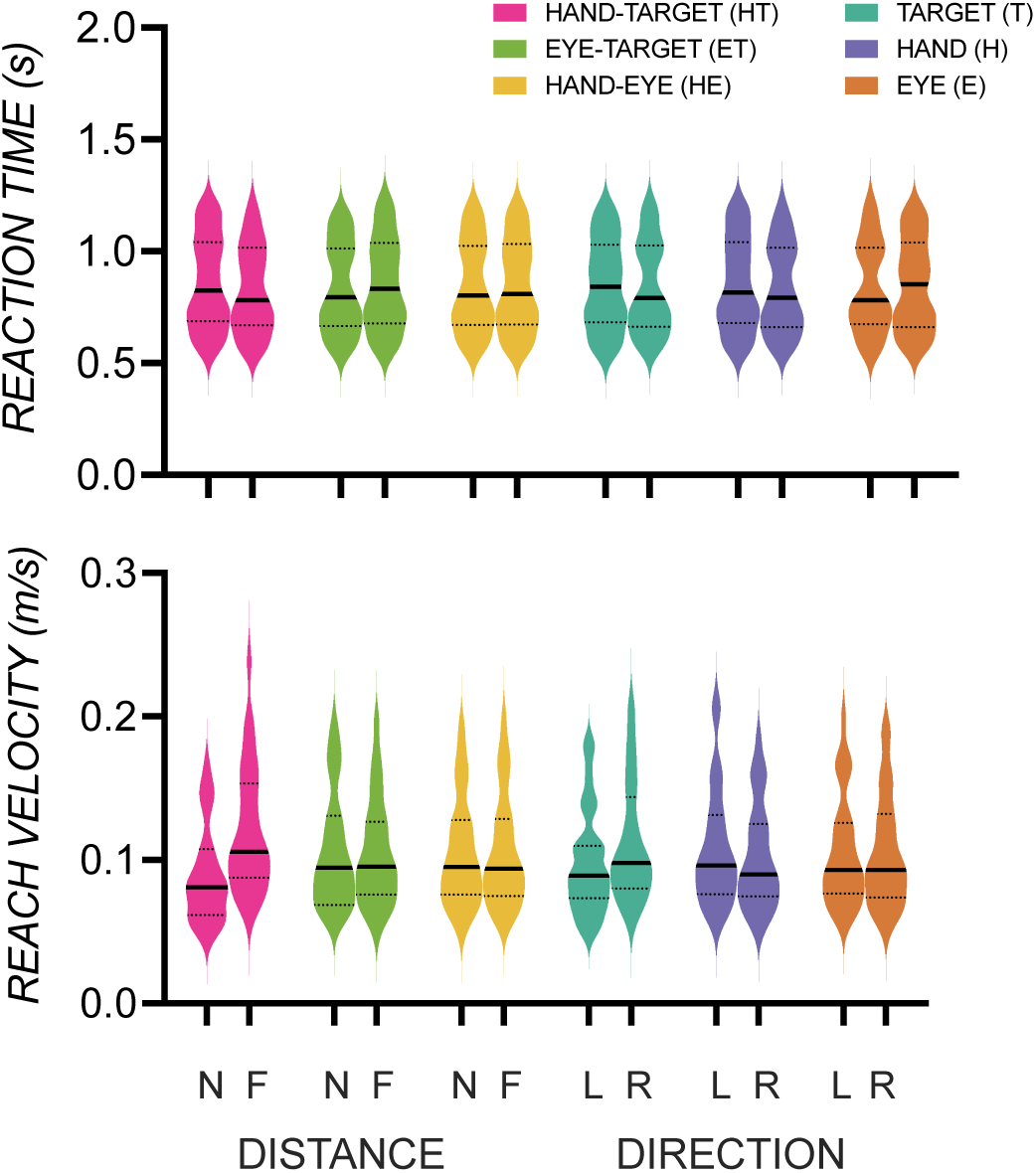
Reaction time (s) (top) and reach velocity (m/s) (bottom) were not significantly different between conditions contrasted in the vRSA of direction [left (L) vs. right (R)] of gaze, hand, and target and distance [near (N) vs. far (F)] between hand and target (HT), gaze and target (ET), and hand and eye (HE).

## Discussion

Motivated by nonhuman primate electrophysiology studies (Batista et al., 1999; Buneo et al., 2002; Batista et al., 2007), we took a task-based fMRI approach exploiting pattern component modeling to determine how distance between target, gaze, and hand position are represented, while also accounting for direction components known to be encoded throughout the human dorsomedial reach pathway. During early planning, rostral PMd and SPL 7a encoded target distance in multiple reference frames while other regions including M1, caudal PMd, and SPL 5m, 5ci, 7pc encoded target distance in a single reference frame that was mostly gaze-centric (except SPL 5m). Notably, all regions encoding distance information also encoded direction information independently. Consistent with our hypothesis, during late planning, distance encoding in multiple reference frames was magnified. In addition, previously widespread direction-based effects were somewhat diminished during late relative to early planning. These results support the hypothesis that the human central nervous system can flexibly define and transform movement-related parameters in multiple reference frames to enable successful goal-direction action.

A strength of the vRSA approach is to test for the independent contribution of each component to an overall spatial activity pattern while accounting for all included components. In this way, an ROI can be sensitive to more than one component. Mixed representations pose a greater challenge for winner-takes-all univariate methods, or standard multivariate pattern decoding algorithms that assume representational singularity within a region (Dubois et al., 2015). Here we combined vRSA with a novel behavioral approach that not only dissociated distance and direction features but minimized between-condition activity pattern differences due to accuracy, speed, or visual acuity. With this approach, we derived strong empirical support for independent representations of distance and direction of an upcoming reach along the dorsomedial reach pathway.

Dorsomedial reach areas encoded a target’s distance, demonstrating that parameters other than direction are specified by recruiting parietal and not just premotor regions (Fabbri et al., 2012). Findings in these regions and across epochs are generally consistent with the hypothesis that distance, like direction, is encoded in multiple reference frames. While we are the first study in humans to isolate and identify gaze– and body-centric distance representations across the dorsomedial reach pathway, independent of direction-based representations, our results align with a series of recent non-human primate studies showing mixed reference frame encoding in the medial sector of posterior parietal cortex, i.e. SPL (Hadjidimitrakis et al., 2014b; Hadjidimitrakis et al., 2014a; Bosco et al., 2016; Piserchia et al., 2016; De Vitis et al., 2019; Hadjidimitrakis et al., 2022). Representations of distance in multiple reference frames suggest a similar circuitry that flexibly transforms between gaze and body-centered coordinate spaces for distance as it does for direction.

Consistent with our hypotheses, distance representations in multiple reference frames were magnified during later planning. During early planning, SPL 7a, PMd rostral, and PMd rostral-sulcal regions represented distance in multiple reference frames. Other regions (SPL 7pc, 5ci, PMd caudal, M1) only represented gaze-centric distance, whereas somatosensory SPL 5m represented target distance in body coordinates. In late planning, all regions with credible evidence represented distance in body and gaze-centered coordinates (SPL 7a, 5m, PMd caudal, rostral, rostral-sulcal, M1). Gaze-centric representations during early planning and body-centric distance representations that were only observed in late planning (e.g., PMd caudal, M1) are consistent with nonhuman primate work showing the encoding of target parameters in gaze-centric preceding body-centric reference frames (Bremner and Andersen, 2012; Cappadocia et al., 2017). Intriguingly, SPL 5m represented the target distance in only a body-centric reference frame in early planning, then both in both body– and gaze-centric reference frames in late planning, suggesting that gaze-centric need not always precede body-centric representations. These results provide empirical support for fluidity in how and when a region represents target distance, with increasing support for spatial overlap in parametric representations of an upcoming target in multiple frames (Boussaoud et al., 1998; Buneo et al., 2002; Batista et al., 2007; Marzocchi et al., 2008; Bernier and Grafton, 2010; Beurze et al., 2010; Chang and Snyder, 2010; Bremner and Andersen, 2012; Hadjidimitrakis et al., 2014a; Leoné et al., 2015; Bosco et al., 2016; Piserchia et al., 2016; Cappadocia et al., 2017; Hadjidimitrakis et al., 2017; De Vitis et al., 2019; Magri et al., 2019; Hadjidimitrakis et al., 2020).

Distance representations in multiple reference frames across motor, premotor, and parietal reach nodes argue against a strict posterior-to-anterior gradient in sensorimotor transformations for skilled action, where posterior areas encode motor goals in gaze-centered reference frames and more anterior regions encode targets in body-centered reference frames. Previous studies have shown that both posterior and anterior regions within parietal and premotor areas integrate motor features in visual and proprioceptive coordinate space (Boussaoud et al., 1998; Buneo et al., 2002; Bernier and Grafton, 2010; Chang and Snyder, 2010; Bremner and Andersen, 2012; Hadjidimitrakis et al., 2014a; Piserchia et al., 2016). Thus, sensorimotor transformations might not be exclusively segregated along a posterior-to-anterior axis. If anything, our results give evidence for gaze-to body-centered transformations occurring temporally across multiple nodes of the dorsomedial reach pathway.

Most regions encoding distance information also encoded direction information independently during early planning, which, consistent with non-human primate work, suggests overlap in neuronal populations that encode direction and distance features critical for an upcoming goal-oriented reach (Messier and Kalaska, 2000; Churchland et al., 2006; Hadjidimitrakis et al., 2014b; Bosco et al., 2016; Piserchia et al., 2016; De Vitis et al., 2019; Hadjidimitrakis et al., 2022). Stronger representational specificity for direction than distance components during early planning, and magnified distance representations in later planning support serial processing accounts showing that distance-related activity evolves more slowly than direction-related activity within premotor (Messier and Kalaska, 2000; Churchland et al., 2006) and parietal regions (Hadjidimitrakis et al., 2014b; Hadjidimitrakis et al., 2014a; Piserchia et al., 2016; Hadjidimitrakis et al., 2022). Critically, our results show that the human dorsomedial reach pathway is organized similarly, in that parameterizing the direction and distance of an upcoming reach action can be achieved by recruiting the same functional regions.

Heightened gaze and body-centered representational specificity of a target’s distance were seen in conjunction with a relative dissipation of direction-based representations during late planning. Many regions representing target positions (SPL 7pc, 7p, 5m, PMd caudal, M1) and hand positions (SPL 7pc, 7p, 5m, PMd rostral) during early planning more weakly encoded direction-based information during late planning. Unlike during early planning, magnified distance-representational specificity marginally exceeded direction-representational specificity across dorsomedial reach nodes. Stronger directional than distance encoding during early planning aligns with seminal PMd studies with non-human primates and more recent studies in parietal regions (Messier and Kalaska, 2000; Hadjidimitrakis et al., 2022). Similarly, transcranial magnetic stimulation in humans of caudal PMd near movement execution disrupts distance but not movement direction (Davare et al., 2015), which aligns with the dissipation of direction representations in caudal PMd observed here. Although direction-based representations relevant to an upcoming reach dissipated in late planning across the frontoparietal network, they were not completely absent. Hand and target position encoding persisted in three out of eight regions (hand: SPL 7a, 5l, M1; target: SPL 7a, 5l, PMd rostral), whereas representations of gaze position were mostly stable and invariant through early and late planning.

Weaker target direction encoding than distance encoding in caudal PMd during late planning are inconsistent with earlier non-human primate studies on direction and distance encoding that showed significant, and dominant, direction effects throughout the plan period e.g., (Messier and Kalaska, 2000; Churchland et al., 2006). Several reasons might account for these inconsistencies. First, these previous studies did not explicitly manipulate or dissociate eye position from hand/body position relative to upcoming reach targets as we have done here. Thus, it is unknown whether the observed neural activity scales with target direction relative to body or gaze coordinates. With our task design, we can interpret activity pattern differences between left and right-sided targets to reflect varying target position representations in absolute space or relative to the hand or body. We show that absolute or body-centric target direction dissipated in later planning in caudal PMd. Second, our task design was specifically set up to compare credible evidence for distance and direction representations that “compete” with each other in the vRSA pattern component modeling framework. More recent monkey neurophysiology work that adopt similar paradigms show stronger distance than direction encoding consistent with our results e.g. (Hadjidimitrakis et al., 2022). Finally, and perhaps most importantly, our use of an ROI-based analysis of multivoxel patterns fundamentally differs from single-neuron or spike-rate histogram methods, where individual neurons or pockets of neurons may still encode for a specific feature (like direction) even as the broader population activity shifts to another feature (like distance). The fMRI vRSA approach captures population-level activity within the entire region, and therefore may give a more comprehensive view of the region’s function and may account for the observed differences between our findings and work with nonhuman primates.

Nevertheless, we should be cautious in assuming that target direction effects are weaker during late-stage planning. For one, gaze position and target direction are necessarily coupled in our paradigm, and the strong gaze representations observed in many regions over time could be tied to the direction of the upcoming movement. It is important to qualify that gaze position representations persisted in caudal PMd (and other regions with target direction dissipation effects). These activity pattern differences between left– and right-sided gaze positions can reflect varying gaze positions in absolute space or relative to the hand, body, or target. In this way, gaze position effects may be partly driven by target direction differences relative to gaze. That all said, target direction representations were not absent, persisting in SPL7a, 5l, and areas PMd rostral areas, in addition to widespread gaze representations. Given the fluidity of these representations over time, it is likely that regions including M1 represent this information after the go-cue for accurate reaching. In line with this, Fabbri and colleagues (2012) showed during movement execution parietal encoding of direction and distance, consistent with what we show here during early and late planning (within 7a surveyed here). Similarly, they showed large to small adaptation in PMd suggesting nuanced amplitude encoding in direction-sensitive premotor areas.

Gaze fixation representations were widespread throughout the dorsomedial reach pathway and credibly evident across planning epochs, with large median Bayes factors. We caution against drawing major inferences relating to components with credible evidence with varying Bayes factor magnitudes, as our design was not set up to compare the size of component effects. Specifically, larger Bayes factor magnitudes for initial gaze fixation likely stem from contrasting activity patterns with greater positional differences between left and right gaze direction on the task board, than direction and distance differences contrasted in other components. It might have been more challenging to detect spatial activity differences between reaching toward left and right targets that are more closely spaced to each other than left and right gaze points. That said, previous fMRI studies have detected spatial activity pattern differences for left and right reach targets using similar or smaller directional and distance differences than that sampled here (Batista et al., 2007; Bernier and Grafton, 2010; Beurze et al., 2010; Fabbri et al., 2010, 2012). The narrow orientation of the hand and target buttons was selected to limit movement, allowing participants to reach from the hand to the target buttons without requiring movement of the entire arm. We also chose target and gaze positions ensuring that targets on all trials would always fall within an observer’s near peripheral zone within which there are little changes in visual acuity (Millodot et al., 1975; Larson and Loschky, 2009). In this way, we minimized between-condition activation pattern differences between near vs. far distances between the gaze and target to be driven by visual acuity differences. Another methodological limitation of the current design is an inability to differentiate between anchor points for body-centered representations since hand, shoulder, and head-centered coordinate systems would be indistinguishable. Finally, with all movements executed along a single, horizontal axis, our results are specific to distance or movement amplitude in 2D (not 3D) space.

Encoding a goal in multiple reference frames supports the flexibility of the brain to convert or switch a target’s distance, as well as its direction, relative to the gaze or body. These motor parameters (direction and distance) of an upcoming goal-directed reach can undergo sensorimotor transformations between gaze– and body-centric frames by recruiting the same areas. This aligns with the hypothesis that such transformations contribute to the brain’s flexibility. Neuronal encoding of motor parameters in multiple coordinate frames, as demonstrated in our study, supports computational models emphasizing the flexibility, efficiency, and power of individual nodes coding diverse inputs to optimally combine or transform extrinsic or intrinsic representations (Körding and Wolpert, 2004; Sober and Sabes, 2005; McGuire and Sabes, 2009).

Brain-computer interface technology decoding motor intention through neural activity for generating movement goals has advanced rapidly with the integration of sensory information (Flesher et al., 2021). Harnessing sensorimotor integration’s benefits requires carefully delineating the timing and reference frame in which multiple motor parameters are specified for goal-directed action. Representations of direction and distance in multiple reference frames observed here are consistent with computational models specifying a flexible brain that can anchor motor parameters to the most reliable or task-applicable sensory source. Future work should clarify how sensorimotor transformations within and between motor parameters unfold with uncertainty in the neural circuitry supporting goal-directed action, with sensorimotor declines present in many neurological disorders at birth (Bleyenheuft and Gordon, 2013; Gupta et al., 2017) and later in life (Abbruzzese and Berardelli, 2003), including healthy aging (Cham et al., 2007; Seidler et al., 2010; Kuehn et al., 2018; Cassady et al., 2020). The present study’s task-based fMRI approach with pattern component modeling contributes to closing a critical divide between nonhuman and human neurophysiology by characterizing how and when goal-relevant representations for upcoming action are encoded.

## Author contributions

Conceptualization, D.B. and M.M., Methodology, D.B., N.D., and M.M., Software, D.B., J.S., and M.M., Formal Analysis, N.D., A.H.C., and M.M., Investigation, A.H.C and M.M., Writing – Original Draft, A.H.C. and MM, Writing – Review and Editing, A.H.C., D.B., N.D., J.S., and M.M., Visualization: A.H.C. and M.M., Supervision: M.M., Funding Acquisition, M.M.

## Acknowledgments

This work was supported by the Wu Tsai Human Performance Alliance and the Joe and Clara Tsai Foundation. We thank David Badcock for his help with visual angle and acuity considerations, Scott T. Grafton, and Giacomo Ariani for their helpful input on data analyses and interpretation. We thank Nicole Stoehr and Suhana Ahamed for assisting with developing and piloting the task design, and Kaden Coulter for assisting with analyses.

